# Strain-dependent differences in the capacity of peste des petits ruminants virus to infect antigen-presenting cells

**DOI:** 10.64898/2025.12.08.692920

**Authors:** Aadel Bouziane, Vincent Lasserre, Roger-Junior Eloiflin, Manon Chambon, Camille Piro-Mégy, Samia Guendouz, Philippe Holzmuller, Philippe Totté, Arnaud Bataille

## Abstract

The ability of morbilliviruses to infect antigen-presenting cells (APCs) plays a major role in infection outcome, because these cells transport peste des petits ruminants virus (PPRV) from the respiratory tract to the lymph nodes before spreading to lymphoid and epithelial tissues. This study used two cell models, — monocyte-derived macrophages (MoMs) and monocyte-derived dendritic cells (MoDCs), — from three host species (goat, sheep and cattle) to evaluate the differential ability of APCs to be infected by two strains of PPRV with contrasting virulence in goats. Infection was analysed using flow cytometry, infectious titers (TCID₅₀/mL), RT-qPCR and immunofluorescence. Goat and sheep MoDCs proved highly permissive to both strains, with viral titers at 48 hpi in the range of 10⁴–10⁵ TCID₅₀/mL and a high percentage of infected cells. Cattle MoDCs were weakly infected and had no detectable viral titer, as expected for this dead-end host. In comparison, MoMs showed reduced permissiveness to PPRV in goats but not in sheep. Post-infection supernatants in goat MoMs infected with the low-virulent strain showed low or undetectable TCID₅₀/mL, despite persistence of detectable viral genome up to 96 hpi. Immunofluorescence revealed double-stranded RNA but no nucleoprotein of the low-virulent strain in goat MoMs, suggesting a partial engagement of the viral cycle, resulting in a restricted or semi-permissive infection. Overall, these results suggest that this combined in vitro model of MoDC and MoM infections would be useful in anticipating host susceptibility and PPRV strain virulence, providing complementary tools for PPR control efforts.

**IMPORTANCE:** Peste des petits ruminants (PPR) disease affects goats, sheep and multiple other hosts across the globe, with major impacts on economies, food security and biodiversity. The sensitivity of host species to PPRV infection varies considerably, with clinical manifestations and disease progression being shaped by the viral strain, host species and breed infected. In vitro models focusing on immune cells, which are the primary target of PPRV could provide a relevant tool for studying the mechanisms of viral adaptation to the host and cellular antiviral responses, while contributing to surveillance and control strategies. The results obtained here suggest that a dual-cell model focusing on macrophages and dendritic cells could be useful for pre-screening studies aimed at assessing the susceptibility of a given host species to a PPRV strain and estimating its relative virulence. Testing this in vitro approach in additional host species and a wider range of strains is required to confirm its relevance.

## INTRODUCTION

Peste des petits ruminants virus (PPRV) is a member of the Paramyxoviridae family, belonging to the *Morbillivirus* genus, which includes the measles virus (MeV), rinderpest virus (RPV) and canine distemper virus (CDV). PPRV has a linear, non-segmented, single-stranded RNA genome with a negative polarity ^1^. First identified in the Ivory Coast in 1942, PPRV has since spread extensively, becoming endemic in large regions of Africa, the Middle East, and Asia ^1,2^, and has recently emerged in several European countries ^3,4^. The virus primarily affects domestic small ruminants, such as goats and sheep, but it can also infect suids and a variety of wild species within the order Artiodactyla, including antelopes and gazelles ^5^.

PPRV is transmitted primarily through direct contact with infected animals, via respiratory secretions, feces, and contaminated feed or water sources. PPR can cause severe clinical manifestations in sheep and goats, with high morbidity and mortality, leading to significant economic losses for farmers. Given its high transmissibility and significant economic impact, PPR is classified as a notifiable disease by the World Organisation for Animal Health (WOAH, formerly OIE) ^6^, and has been the focus of a control programme jointly led by WOAH and the Food and Agriculture Organization of the United Nations (FAO) since 2015, aiming for its eradication by 2030 ^7^. However, the sensitivity of the host species to PPRV infection varies considerably, with clinical manifestations and disease progression being shaped by the viral strain, host species and breed infected^8,9^. A better understanding of PPRV virulence and host susceptibility is considered a research priority for disease control ^10^.

The main targets of PPRV and other morbilliviruses are immune cells via the SLAMF1 (Signalling Lymphocytic Activation Molecule Family 1) receptor, also known as CD150 ^11,12^, and epithelial cells via the Nectin-4 receptor ^13^. At the early stage of infection, morbilliviruses infect antigen-presenting cells (APCs), such as macrophages and dendritic cells, which transport the viral particles to lymphoid organs, in particular lymph nodes and tonsils, where lymphocytes reside ^1^. Infection of epithelial cells occurs once viraemia is established, with the Nectin-4 receptor playing an essential role in viral dissemination and excretion ^11,13^.

For PPRV, direct infection of APCs in vivo has not yet been demonstrated. However, draining lymphoid tissues such as the tonsils and lymph nodes, are the first major sites of replication ^14^. Recent studies have shown that monocyte-derived dendritic cells are permissive to PPRV in vitro ^15^. Permissiveness is defined as the ability of the virus to replicate and produce infectious virions. We know that for other morbilliviruses permissiveness varies depending on the cell type and the virus infecting these cells ^16,17^. Due to the importance of APCs in morbillivirus infection, it is possible that the observed variability in PPRV virulence and host susceptibility can be related to differences in the viral replication capacity in APCs. Among APCs, dendritic cells and macrophages are essential for detecting viral infections and initiating innate immunity^18,19^.

So, the study of infection of monocyte-derived macrophages (MoMs) and monocyte-derived dendritic cells (MoDCs) could be a relevant and reproducible model for comparing the permissiveness of host species to PPRV. Here, we will focus on the infection of MoDCs and MoMs from goats, sheep and cattle (the latter being identified as dead-end hosts for PPRV ^20^) with two strains of PPRV. The two strains used in this study, Morocco 2008 (MA08) and Ivory Coast 1989 (IC89), demonstrated different levels of virulence in certain goat breeds ^21^. Strain MA08 belongs to lineage IV of PPRV and was isolated from an Alpine goat that was severely affected during the 2008 outbreak in Morocco ^22^. The IC89 strain belongs to lineage I. It was isolated from a goat in Ivory Coast in 1989 and is considered to be a mildly virulent strain, inducing a mild disease in various goat breeds ^21,23,24^, although it is used as a virulent strain for the study of immune response in sheep ^25^.

Based on representative known virulence to goats of the two strains of PPRV, low for IC89 and high for MA08, the aim of this study was to assess the comparative in vitro replication of these PPRV strains in both MoMs and MoDCs of goats, sheep and cattle. The research question was to evaluate whether any difference could be observed between these APCs in relation to PPRV strain virulence.

## RESULTS

### 1. PPRV infections of MoMs and MoDCs

To determine the infectivity of the two PPRV strains, IC89 and MA08, for macrophages and dendritic cells, circulating monocytes from the blood of the different animal species studied were differentiated into dendritic cells (MoDCs) and macrophages (MoMs). These two cell types were infected with each of the viral strains studied, IC89 and MA08, for 48 hours at a multiplicity of infection (MOI) of 0.1. Infection was assessed by flow cytometry after intracellular labelling with an antibody targeting the virus nucleoprotein (anti-NPPRV) coupled to a fluorochrome.

In goat MoMs, an average of 20% of the cells were positive for the N protein of PPRV strain IC89, compared with an average of 32.6% for those infected with strain MA08. However, no significant difference was observed (Figure 1A). In addition, for goat MoDCs, no significant difference was observed between the two viral strains and in most cases the cells were positive for NPPRV with a range of 40 to 73% (Figure 1B). Intercellular comparison (i.e. MoMs versus MoDCs) revealed a significant difference in the rate of NPPRV-positive cells, with MoDCs being twice as infected as MoMs. This was observed for both PPRV strains (Fig. 2A and 2B).

**Figure 1:**
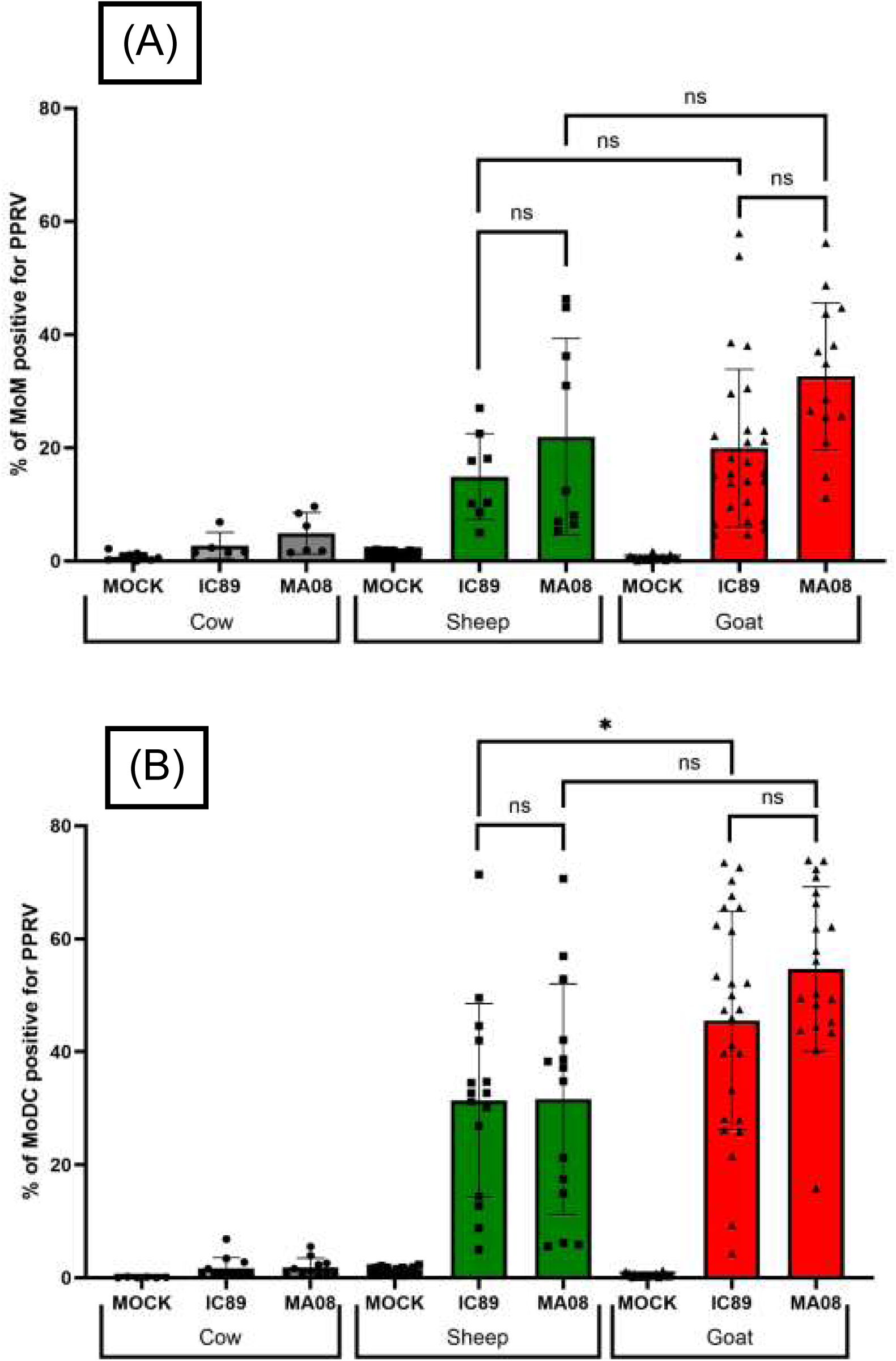
Percentage of MoM (A) and MoDC (B) from cow, sheep and goat, positive for PPRV at 48hpi after in vitro infection with IC89 or MA08 strain. The MOCK group corresponds to uninfected control cells. Data are reported as individual biological replicates from minimum 3 animals per species and per condition with mean ± SD; ns=non-significant; *P<0.05; Statistical analysis was performed using a generalized linear mixed-effects model (GLMM). Post-hoc comparisons were conducted using estimated marginal means with Tukey adjustment for multiple comparisons.

**Figure 2:**
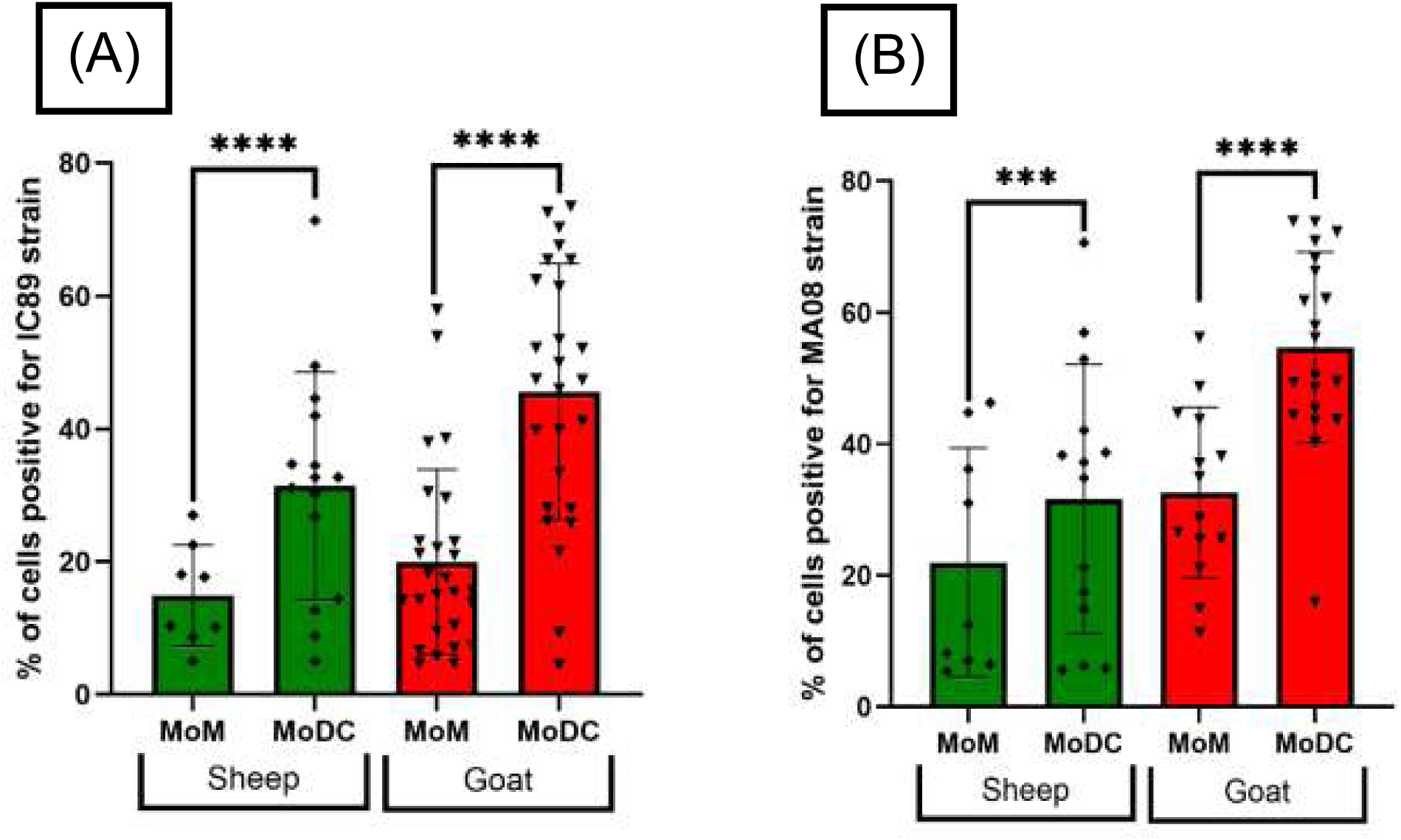
Percentage of MoM and MoDC from sheep and goat, positive for PPRV at 48hpi after in vitro infection with IC89 (A) or MA08 (B) strain. The MOCK group corresponds to uninfected control cells. Data are reported as individual biological replicates from minimum 3 animals per species and per condition with mean ± SD; ***P<0.001; ****P<0.0001; Statistical analysis was performed using a generalized linear mixed-effects model (GLMM). Post-hoc comparisons were conducted using estimated marginal means with Tukey adjustment for multiple comparisons.

For sheep MoMs, there was no significant difference in the rate of NPPRV-positive cells between the two PPRV strains, and the variability of responses was high, with an average rate of 14.7% for IC89 and 21% for MA08 (Fig. 1A). Similarly, for sheep MoDCs, there was no significant difference between these two viral strains in terms of percentage of NPPRV-positive cells. In both cases, the average was 30%, with comparable extreme values (Fig. 1B). A significant difference was observed between sheep MoMs and MoDCs infected with IC89, again MoDCs being about twice as infected as MoMs (P<0.0001) (Fig. 2A). In the case of PPRV strain MA08, the infection rate of MoDCs was significantly higher than in MoMs (P<0.001), with an average of 30% compared to 22% for MoMs (Fig. 2B).

MoMs and MoDCs from cattle were also infected with IC89 and MA08 strains using the same parameters. Flow cytometry analysis after intracellular staining of the NPPRV protein revealed very low levels of NPPRV-positive cells, no matter which cell type and viral strain was used (Fig. 1A, 1B)

### 2. Replication of PPRV in MoMs and MoDCs

To study viral replication in MoMs and MoDCs of different hosts, post-infection supernatants were added to cultures of SLAM-expressing Vero cells (VDS) ^26^ at dilutions from 10^-1^ to 10^-7^ with a minimum of four replicates per dilution. The cytopathic effect (CPE) was observed under the microscope and viral titers were calculated as described in Materials and Methods.

In goats, post-infection supernatants from MoDCs had, on average for both PPRV strains MA08 and IC89, significantly higher titers than supernatants from MoMs (P<0.0001) (Fig. 3A and 3B). For MoMs, the viral titers of the post-infection supernatants for MA08 reached an average TCID_50_/mL of 3x10^3^ (Fig3B) whereas, for the IC89 strain, the viral titer was barely detectable with an average TCID_50_/mL of 34 (Fig 3A) including 9 out of 17 replicates where no titer was detected (Circled in Fig 3A).

**Figure 3:**
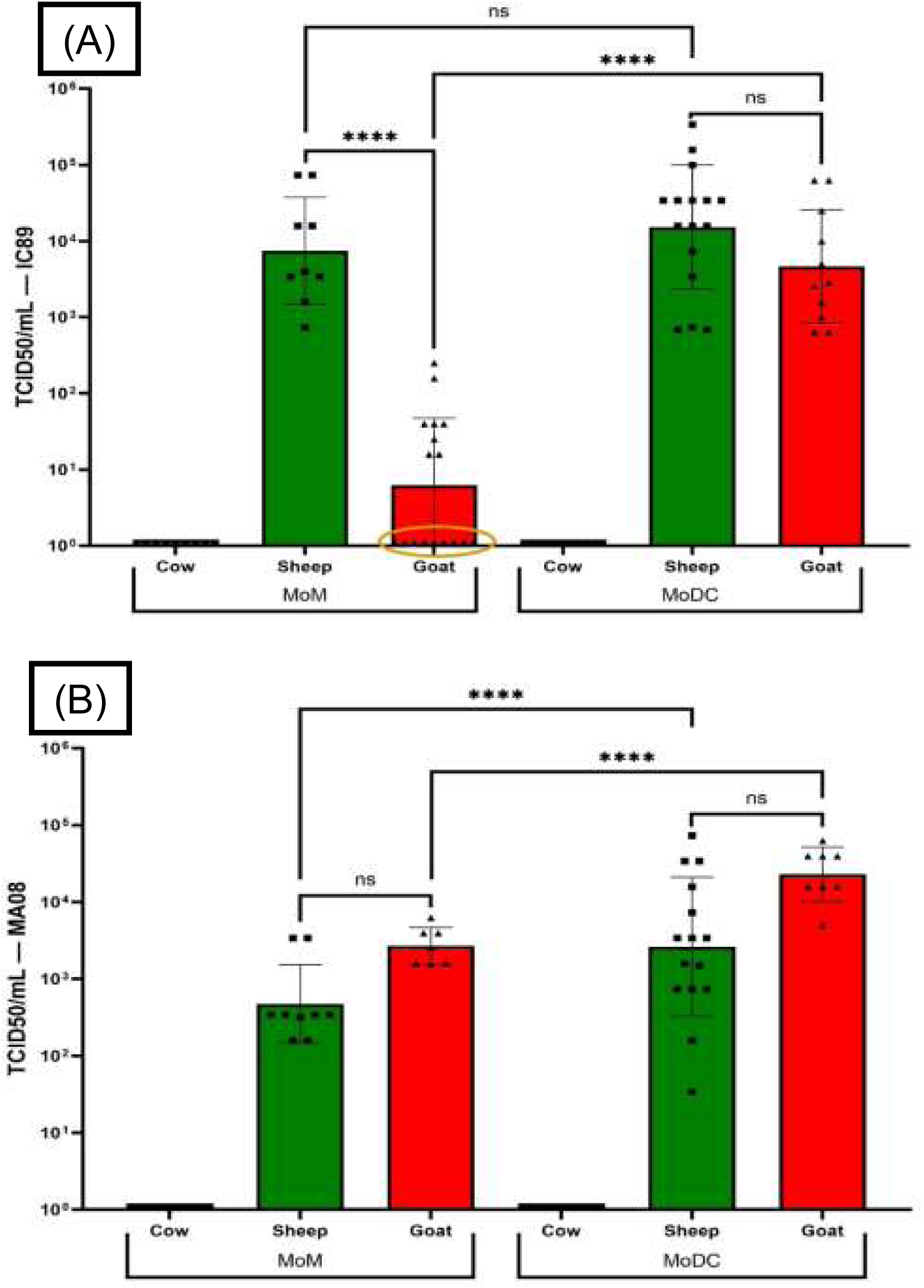
Viral titers (TCID50/mL) of IC89 (A) and MA08 (B) at 48 hpi in cow, sheep and goat MoMs and MoDCs. Nine of seventeen replicates were below the limit of detection in goat MoMs infected with IC89 (circled in A). Data are shown as individual biological replicates from a minimum of three animals per species and per condition with mean ± SD; ns=non-significant; ****P<0.0001. Replicates from goat with no titer observed are highlighted with a circle. Statistical analysis was performed on viral titers using a generalized linear mixed-effects model (GLMM) fitted with a negative binomial distribution. Post-hoc comparisons were performed using estimated marginal means with Tukey adjustment for multiple comparisons

In sheep, post-infection supernatants from MoDCs infected with strain MA08 had, on average, significantly higher titers than those from MoMs infected with MA08 (Fig. 3B) with a TCID_50_ of around 1.2x10^4^ for MoDCs infected with MA08 compared with 9.8x10^2^ for MoMs infected with MA08 (P<0.0001). There were no significant differences between MoMs and MoDCs infected with the IC89 strain even though there was a trend towards higher titers for post-infection supernatants from MoDCs. The IC89 strain yielded a significantly higher mean titer than the MA08 strain in both MoDCs (P<0.001) and MoMs (P<0.0001) (Supplemental fig. S1).

The titration of 48hpi supernatants of the IC89 strain on sheep MoMs versus goat MoMs, indicated a significant difference (P<0.0001), with sheep MoMs exhibiting high titers with an average TCID_50_ of 2x10^4^. In contrast, whereas goat MoMs infected with IC89 had titers that are barely or not at all measurable (Fig. 3A). In cattle APCs, no viral titer was detected in the post-infection supernatants (Fig. 3A and 3B).

### 3. Kinetics of infection of goat MoMs by PPRV IC89 strain

#### a. Infection and replication

After observing low or non-detectable viral titers in supernatants of goat MoMs infected for 48 hours with the IC89 strain, infection kinetics were studied to see if replication was somewhat delayed for this low virulent strain. Goat MoMs were infected for 24 h, 48 h, 72 h and 96 h, at an MOI of 0.1. As previously, the rate of NPPRV-positive cells was measured using a flow cytometer and post-infection supernatants were collected and viral titers were determined.

The results show that the rate of infected cells is relatively stable over time, with an average NPPRV positivity of around 30% regardless of the infection time. No significant increase in the percentage of NPPRV-positive cells was observed between 24hpi and 96hpi (Fig. 4A).

**Figure 4:**
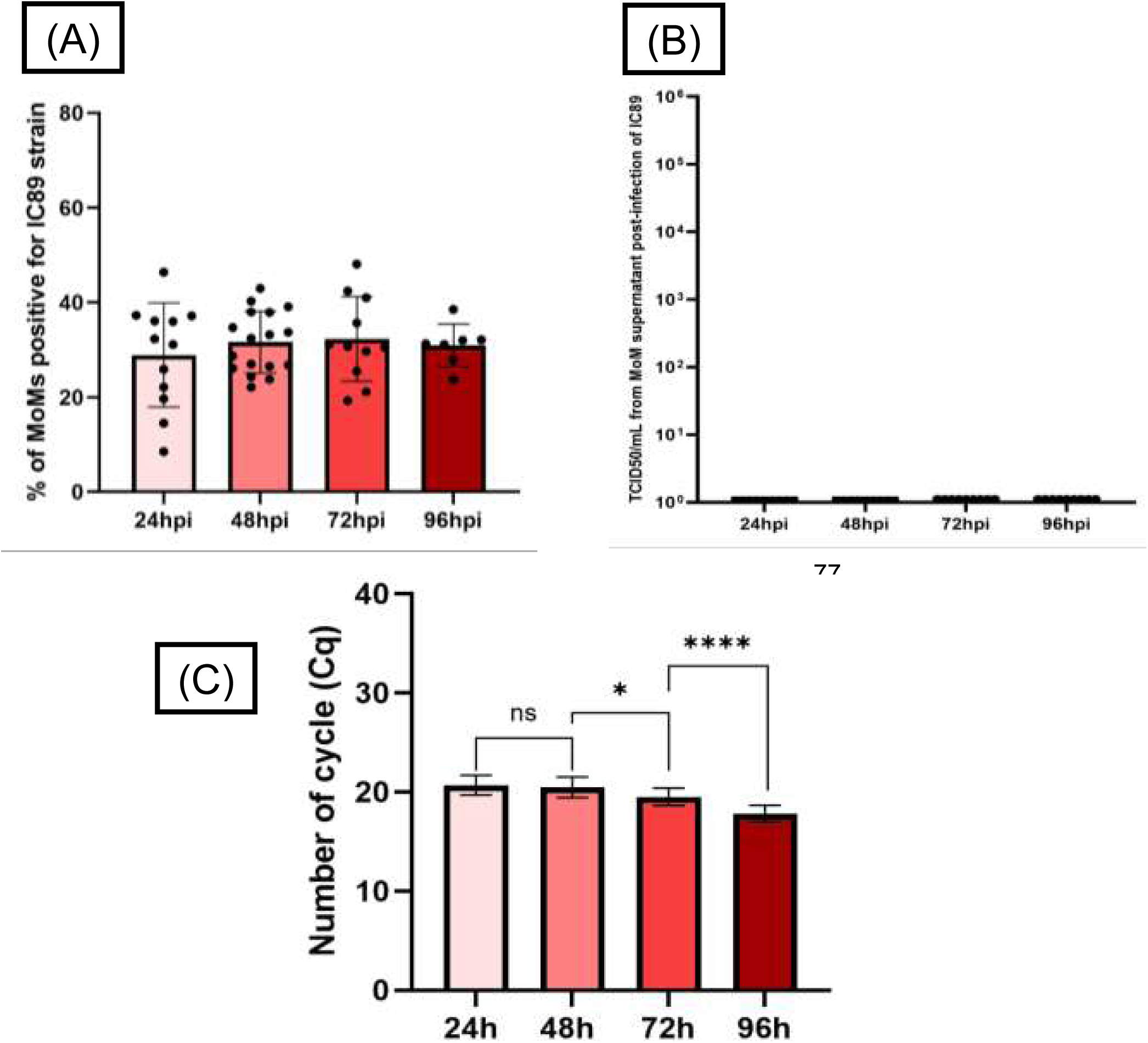
(A) Kinetics of infection of goat MoMs by strain IC89 of PPRV. Percentages of NPPRV+ cells were determined by flow cytometry at 24, 48, 72, and 96hpi. Data are shown as individual biological replicates with 3 goats and with minimum n=7 replicates, mean ± SD. A beta-binomial GLMM revealed no significant effect of time post-infection on infection rates (all P > 0.76). Post-hoc pairwise comparisons using Tukey-adjusted estimated marginal means confirmed the absence of significant differences between 24, 48, 72 and 96 hpi (all adjusted P > 0.99). **(B) Viral titration kinetics of MoMs supernatants from goats infected with strain IC89 at 24, 48, 72 and 96hpi.** Data are shown as individual biological replicates from 3 goats with n=9 replicates per time point. **(C) Kinetics of viral shedding in MoMs supernatant infected by IC89 strain at 24h, 48h, 72h and 96h.Viral shedding was monitored by RT-qPCR (expressed in Cq) detecting the presence of the PPRV N gene.** Data are shown as individual biological replicates from 3 goats with n ranging from 12 to 20 replicates per time point, mean ± SD; *P<0.05; P<0.0001. Statistical analysis was performed using a linear mixed-effects model (LMM) fitted by restricted maximum likelihood (REML). Pairwise comparisons were conducted using estimated marginal means with Tukey adjustment for multiple comparisons (Kenward–Roger approximation for degrees of freedom).

The post-infection supernatants from IC89-infected goat MoMs were collected in triplicates per animal throughout the kinetics, and titrated on VDS. No TCID_50_/mL viral titers could be detected for any time point during the kinetics, with values remaining below the test detection threshold (Fig. 4B), and cytopathic effects (CPEs) sometimes observed in a single well of the first dilution (10^-1^). The titration plates used for this kinetic analysis were then labelled with the monoclonal antibody 38.4 targeting the NPPRV coupled with tetramethylrhodamine isothiocyanate (TRITC) in order to observe the presence or absence of the virus at the cellular level. We observed the presence of the virus in VDS cells in the first dilution, and sometimes up to the second and third dilutions (10^-2^ and 10^-3^). However, this detection occurred in the absence of a cytopathic effect (CPE), or with CPE limited to a single well in the first dilution, which is insufficient to calculate a reliable viral titer (Supplemental fig. S2).

#### b. Detection of viral RNA

To determine the presence of viral genomic RNA, an RT-qPCR was performed on post-infection supernatants throughout the kinetic. The cycle number (Cq) between 24 hpi and 48hpi remained stable at around 20.5. A decrease in the Cq value was first observed at 72hpi, falling to an average of 19.5, with an even more significant decrease at 96hpi, when the Cq averaged 17.8 (Fig. 4C), representing an increase in genetic material of the IC89 strain detected in supernatants.

### 4. Detection of dsRNA and NPPRV protein in goat MoMs and MoDCs

To investigate a possible blockage or slowdown in the replication cycle of the IC89 strain in goat MoMs, immunofluorescence was used to detect separately double-stranded RNA (dsRNA), an indicator of the accumulation of aberrant viral RNAs in morbilliviruses, and the PPRV nucleoprotein (NPPRV), a marker of viral protein expression, in infected MoMs. The profiles obtained were compared with those observed in goat MoDCs. A positive dsRNA signal was observed in both cell types in the presence of IC89, although the dsRNA signal appeared slightly less intense in MoMs (Fig. 5A). The dsRNA signal appeared stronger in goat MoMs when infected by MA08 (Supplemental fig. S3). NPPRV is detected in goat MoDCs infected with IC89, but not in goat MoMs (Fig. 5B).

**Figure 5:**
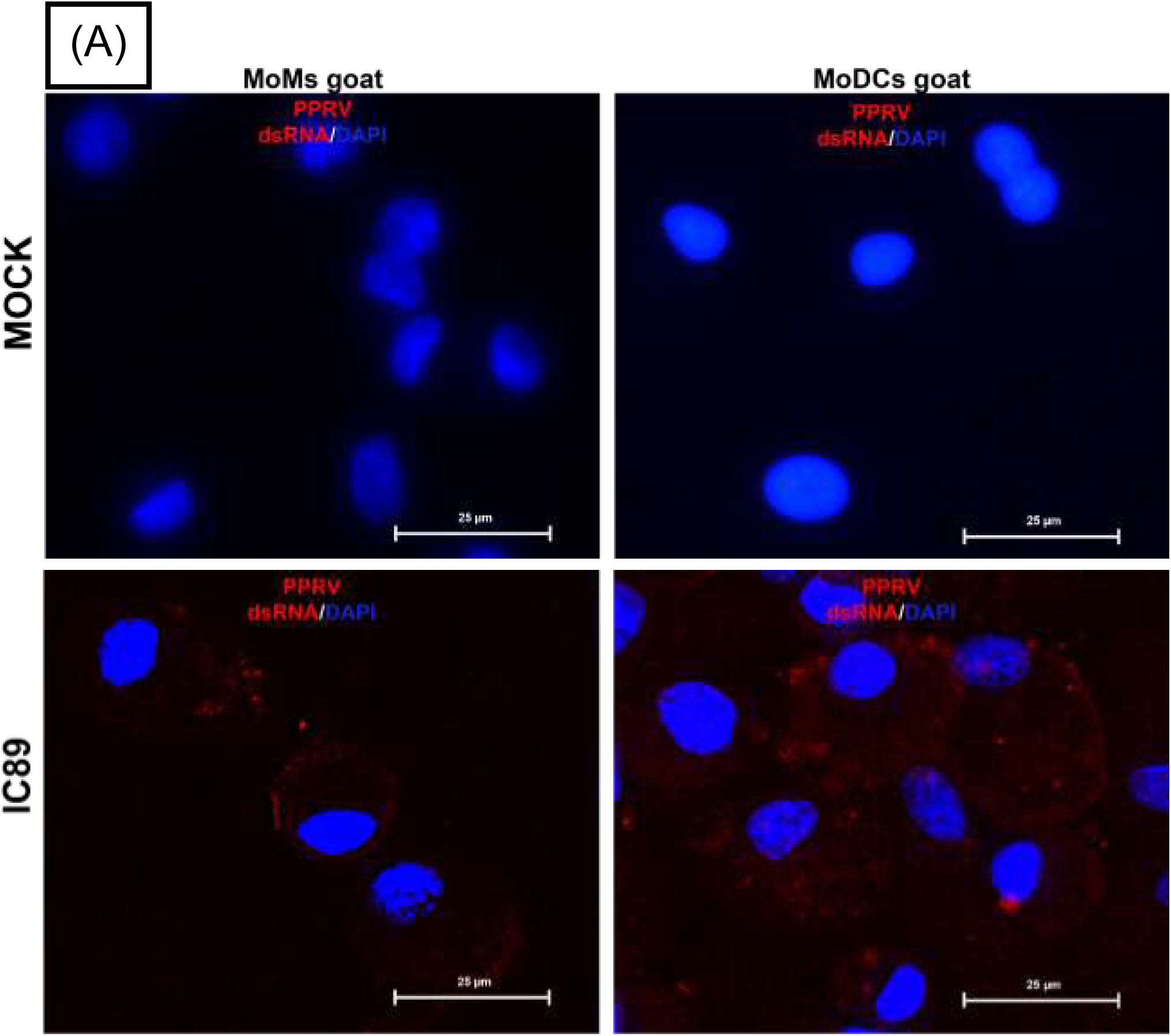
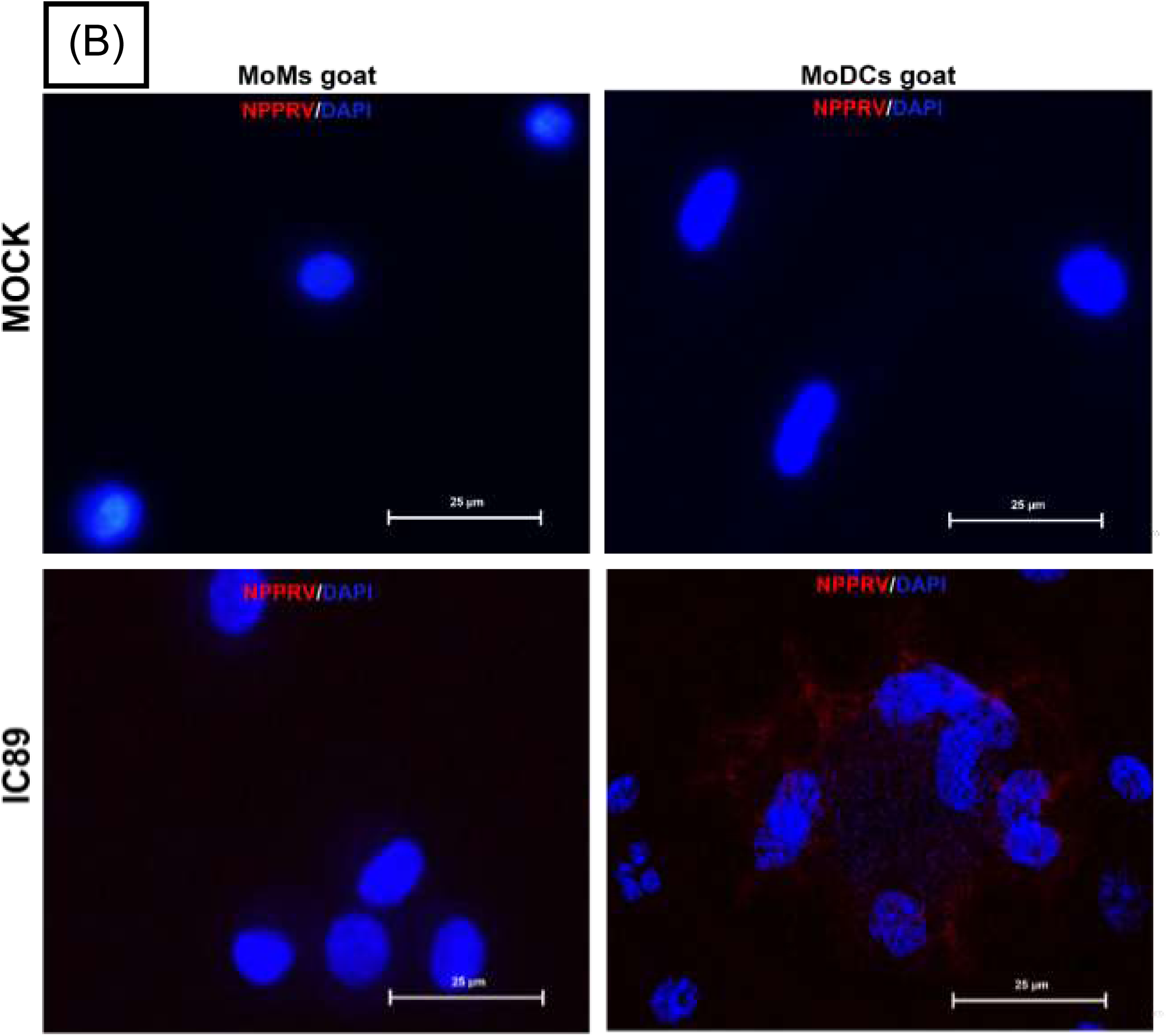
Immunofluorescent detection of viral double-stranded RNA (dsRNA) (A) and Nucleoprotein PPRV protein (NPPRV) (B) in goat MoMs and MoDCs infected with IC89. Infected and uninfected (MOCK) MoMs and MoDCs were stained with DAPI to identify cell nuclei, dsRNA (A) and NPPRV (B) are detected here in red, scale bar = 25 µm.

## DISCUSSION

PPRV infection can lead to a variety of outcomes ranging from acute symptoms and death of the infected host to subclinical infection. Surveillance and control measures must be adapted to provide an efficient response. However, we have limited capacity to predict disease outcomes in different hosts and prepare accordingly. A better understanding of differences in sensitivity of the host to viral strains, studied in vitro, would make it possible to anticipate disease impacts and develop a predictive model that can demonstrate not only the sensitivity of a host to different PPRV strains but also the differences in virulence expressed by the same strain depending on the host ^27,28^. To move towards this objective, we selected two strains of PPRV with distinct virulence profiles in goats, mildly virulent IC89 and highly virulent MA08 ^24,29,30^. Goat and sheep cells were used because these species are the main natural hosts of PPRV, while bovine cells served as a relevant negative control, as cattle are considered a dead-end host ^20,31^.

This in vitro experimental model focused on a comparative overview of PPRV infection in MoMs and MoDCs in order to describe possible differences in sensitivity and replication profile between cell types, host species and depending on the viral strain.

Overall, our study shows that MoDCs from goats and sheep are more permissive to PPRV than MoMs. On average, the rate of infected cells (or NPPRV-positive cells) and viral titers are higher for MoDCs than for MoMs. In cattle, the absence of infection and of measurable replication in MoDCs and MoMs confirms the dead-end host status of this species.

Despite significant inter-individual variability observed in the number of cells infected during the experiment, the measurement of viral replication via titration provided a robust and biologically meaningful reading for defining permissiveness. The data obtained 48 hours post-infection at MOI of 0.1 likely reflect the combined effect of several infectious cycles, incorporating both the efficiency of viral replication and the virus’s capacity for secondary spread in cell cultures. Experiments at lower MOI and at different time post infection may give additional information on the PPRV primary infectious cycle in both cells, however our results show that the conditions used in our in vitro experiments are suitable to detect differences of interest for the overall goal of this study.

No significant differences in viral titers were observed between the IC89 and MA08 viral strains in goat MoDCs. However, in sheep MoDCs, the mean titer of the IC89 strain was approximately 0.6 log_10_ higher that of the MA08 strain. MoDCs results concur with the multiple field and experimental studies showing that cattle are not affected and cannot transmit PPRV, unlike the main small ruminant hosts ^20,32^. In small ruminants, MoDCs confirmed their susceptibility to PPRV infection, but did not reveal any significant difference between goats and sheep. This cell model could therefore potentially serve as an indicator of the ability of the virus to infect a given species.

In contrast to MoDCs, MoMs are less permissive but more discriminating host cells for PPRV. In goats, strain MA08 infected MoMs more efficiently than strain IC89. Oppositely, IC89 produced detectable titers that were significantly higher than MA08 in sheep MoMs. Furthermore, the titers observed with IC89 in sheep MoMs were significantly higher than those obtained in goat MoMs. This suggests that the ability of this strain to replicate will depend on the host species and the type of cell infected. Based on this, and on knowledge of the virulence of these strains in vivo ^29,21,24,30,33^, we propose that this infection model with MoMs could serve as an indicator of strain-specific virulence in small ruminant breeds and may highlight host-specific signatures of viral adaptation.

For the remainder of the study, infection kinetics were performed on goat MoMs infected with IC89, since the discrepancy was most pronounced in this model (i.e. cellular infection was detected by flow cytometry, but the viral titer was undetectable). The objective was to determine whether the absence of a viral titer was due to a delay in replication or a persistent blockage of the viral cycle. Throughout the kinetics (24h to 96h infection), the rate of infected cells remained stable, averaging between 28 and 32% with no significant increase, suggesting continuous infection of a sub-population of macrophages present in the culture. Titrations of supernatants at all time points remained negative, indicating the absence of measurable infectious virion production. Nevertheless, RT-qPCR revealed a significant decrease in Cq between 48, 72 and 96 hpi, reflecting a gradual accumulation of extracellular viral genomic RNA. It should be noted that RT-qPCR does not distinguish between infectious and non-infectious viral particles. In addition, immunofluorescence microscopy of the titration plates showed that viral antigens were present in the cells because of the NPPR protein detection in VDS cells, at different levels but without CPE, which supports the viral titer determined in supernatants and the cytometry results.

Despite the absence of measurable infectious titer, the detection of viral RNA in the supernatants suggests an incomplete or an abortive replication cycle. This could result in the release of non-infectious or defective viral particles, which has already been observed with several RNA viruses^34–36^, including morbilliviruses^37–40^.

In negative-sense single-stranded RNA viruses, such as PPRV, viral genome replication is closely linked to nucleocapsid assembly, meaning that dsRNA is not considered a reliable indicator of productive replication ^41^. Nevertheless, the presence of dsRNA is more likely to be an indicator of the accumulation of aberrant viral RNA, such as defective interfering RNA (DIs) or defective viral genomes (DVG)^39^. Such DIs and DVGs can be produced during virus production, but the contrasting levels of dsRNA observed in MoMs and MoDCs infected with IC89 suggests differences in the accumulation of these particles during infection. Importantly, no NPPRV signal was detected in MoMs infected with the IC89 strain, unlike in MoDCs. This would suggest that the IC89 viral cycle in MoMs appears to be engaged until the accumulation of aberrant viral RNA forms (DIs or DVGs), while nucleoprotein expression or accumulation is severely limited or absent. Therefore, while partial expression of the viral genome is detectable, this does not result in the production of infectious particles, indicating a restricted or semi-permissive infection of goat MoMs by the IC89 strain.

Such a restriction may be associated with the activation of interferon-dependent antiviral pathways known to limit viral protein synthesis^42,43^.

Further work is needed to precisely understand the observations made within this promising model in order to evaluate its usefulness. It would be interesting to move towards a multiomics approach to understand the different mechanisms involved in PPRV infection of MoMs and MoDCs, in particular those related to innate antiviral immune activity and the link between different immune signalling pathways and the non-structural proteins C and V of morbilliviruses. It would also be relevant to quantify the different production of cytokines in the post-infection supernatants of MoMs and MoDCs, and to perform proteomic analysis of these supernatants to detect and characterise viral proteins released during infection. Finally, observing the state of the virion as it leaves the cells under electron microscope would provide additional structural information on the viral particles produced and their infectious capacity.

In summary, here we have highlighted the difference in permissiveness to PPR virus strains between MoDCs and MoMs. As demonstrated by the results we obtained with the IC89 and MA08 strains in the two cell models obtained from three different hosts (goats, sheep and cattle), in vitro infections of MoDCs may enable us to determine whether a given host is susceptible to the PPR virus, and MoMs could serve as a cell model to determine the virulence of a given strain among susceptible hosts. With the IC89 strain, an abortive infection is observed in goat MoMs, while sheep MoMs are completely permissive. MoMs from goat or sheep also remain permissive for the MA08 strain. The use of bovine MoMs and MoDCs in this study constitutes a relevant negative control due to its status as a dead-end host. This combined model of two cell types (MoDCs and MoMs) would be useful for pre-screening studies to assess the sensitivity of species to a given strain of PPRV and thus estimate relative virulence. Testing other PPRV strains with goat, sheep and cattle MoMs and MoDCs, and extending such tests to other host species will confirm the utility of this model.

Since 2024, Europe has been facing the emergence of PPR, with small ruminants infected in seven different countries and showing a wide variety of symptoms ^44–47^. This recent dynamic in the spread of PPRV shows that the epidemiological situation remains unstable and that the virus still crosses borders, making it essential to strengthen risk assessment tools and early detection capacities. The risk is also present in areas that interface with wildlife, with cases observed in atypical hosts and endangered wild ruminants in contact with domestic animals ^48–52^. Furthermore, the impact of PPRV on wild species remains poorly understood, as does their potential role in the transmission or maintenance of the virus and modulation of virulence. As global eradication efforts led by FAO and WOAH intensify, having tools to anticipate host susceptibility and strain virulence would be a major asset in guiding early detection and rapid response measures.

## MATERIALS AND METHODS

### Cell lines and viral production

All viral strains were produced from Vero expressing the canine SLAM receptor (or Vero-Dog-SLAM cells, VDS cells) ^26^. VDS cell were cultured in complete Dulbecco’s Modified Eagle’s Medium (DMEM, Gibco) supplemented with 10% foetal bovine serum (FBS, Dominique Dutscher). Virus stocks were harvested from infected cell supernatants, clarified by centrifugation, and stored at -80°C until use. The PPRV strains IC89 (lineage I) and MA08 (lineage IV) are from productions used in previous experimental studies ^21^. Any risk of genetic drift of PPRV strains during cell passages was assessed by sequencing the full genome of the virus production used for the infection experiments using a protocol described elsewhere^53^. Genomic comparison showed that no mutations were detected following the passages required for production.

Differentiation of macrophages and dendritic cells from CD14+ monocytes Peripheral blood samples were obtained from clinically healthy goats (*Capra hircus*, N = 3), sheep (*Ovis aries*, N = 3) and cattle (*Bos taurus*, N = 6) housed at the CIRAD animal facility (Montpellier, France). Animal care and handling followed institutional ethical guidelines and was approved by the Languedoc-Roussillon regional ethics committee (French CE-LR#36) in an authorised project using animals for scientific purposes APAFIS#2628.

Peripheral blood mononuclear cells (PBMCs) were isolated from the whole blood of the animals studied by Ficoll-Hypaque density gradient centrifugation using Histopaque-1077 (Sigma-Aldrich), following standard protocols ^54^. After collection, viable PBMCs were counted in a hemocytometer using the Trypan Blue exclusion method.

CD14⁺ monocytes were positively selected using CD14 MicroBeads UltraPure (Miltenyi Biotec) and LS MACS columns, according to the manufacturer’s instructions. After elution, CD14⁺ monocytes were cultured in Roswell Park Memorial Institute medium and GlutaMAX^TM^ (RPMI 1640 + GlutaMAX^TM^, Gibco^TM^) supplemented with 10% FBS and 1% antibiotics (Penicillin-Streptomycin). A minimum purity of 80% CD14⁺ monocytes is required to ensure homogeneous differentiation into MoMs or MoDCs (Supplemental fig. S4).

For differentiation into monocyte-derived dendritic cells (MoDCs), cultures were supplemented with in-house bovine recombinant IL-4 and GM-CSF (50 ng/ml) From plasmid prepared with EndoFree Plasmid Purification Maxi Kit Qiagen (reference 12362), cytokines were then produced by transfection on HEK 293T cells. For the differentiation into monocyte-derived macrophages (MoMs), no exogenous growth factors were added and monocytes were allowed to spontaneously differentiate into macrophages under standard culture conditions. This approach, although less controlled than cytokine-supplemented protocols, mimics a more physiological environment and results in a heterogeneous macrophage population^55,56^. Differentiation was carried out for six to seven days with a medium change on day two or three (Supplemental fig. S5). In goats, characterization based on CD205 marker expression was performed to distinguish monocytes from cells differentiated into MoMs or MoDCs (Supplemental fig. S6A). Similarly, the CD209 marker was used in cattle^57^ (Supplemental fig. S6B). To distinguish MoMs from MoDCs in goats and sheep, the CD172a marker was used (Supplemental fig. S6C). In addition, SLAMF9 receptor (homolog of SLAMF1) labeling was also performed on goat cells (Supplemental fig S6D). Viral infection assays were performed at the end of the differentiation period. In addition to phenotyping, microscopic observation (Supplemental fig. S5) confirmed that the cells had been differentiated in agreement with previous work^15,57,58^.

### Viral infection

MoDCs and MoMs were infected with either MA08 or IC89 strains at a multiplicity of infection (MOI) of 0.1. The cells were incubated with virus for one hour at 37°C in serum-free RPMI. After incubation, cells were washed with PBS and cultured for 48 hours in RPMI supplemented with 2% FBS. 48 hours post-infection, the culture supernatants were harvested, aliquoted, and stored at –80°C for downstream analysis.

### Flow cytometry

Following infection, adherent cells in culture were detached using TrypLE^TM^ Express Enzyme 1X (Gibco) at 37°C for 10 minutes. Cells were harvested into PBS, centrifuged at 300 x g) for 5 minutes at 4°C, and fixed with 4% paraformaldehyde (or 4% PFA) for 10 minutes at 4°C.

For intracellular PPRV detection, cells were permeabilized with 0.3% saponin and stained with FITC-conjugated monoclonal antibody (clone 38.4) targeting the N nucleoprotein of PPRV developed by G. Libeau ^59^ and produced by ProteoGenix (Schiltigheim, France). No secondary antibody was required due to conjugation of Mab 38.4 with FITC. Stained cells were analyzed using a BD FACSCanto II flow-cytometer (BD Biosciences) and FlowJo_V10 software (Supplemental fig. S7).

For phenotyping, the primary antibodies anti-CD14 (CAM36A), anti-CD205 (ILA114A), anti-CD209 (209MD26A), anti-CD172a (DH59B) and anti-SLAMF9 CACT206A were provided by WSU Monoclonal Antibody Center Veterinary Microbiology and Pathology (Washington State University – Pullman). The secondary antibodies used, rat anti-mouse IgG1, eBioscience^TM^, PE-Cyanine7 Clone M1-14D12 (Reference: 24-4015-82), Invitrogen by Thermo Fisher Scientific and Alexa Fluor^TM^ goat anti-mouse IgG2a (Reference: A21241), Invitrogen by Thermo Fisher Scientific.

### Titration of PPRV

Viral production and cell culture supernatants were titrated on VDS cells seeded in 96-well plates using the Spearman-Kärber method ^60–63^. Infectious titers were calculated as TCID_50_/mL using the TCID_50_ calculator developed by Marco Binder. (v2.1—20-01-2017_MB* by Marco Binder; adapted @ TWC https://www.klinikum.uni-heidelberg.de/fileadmin/inst_hygiene/molekulare_virologie/Downloads/TCID50_calculator_v2_17-01-20_MB.xlsx). The titration plates were marked with antibody 38:4 (anti-NPPRV) coupled with TRITC in order to observe the presence or absence of the virus (Supplemental fig. S2), according to the Eloiflin *et al.* ^64^ protocol.

### RNA extraction from supernatant and real-time qPCR analyses

Viral RNA was extracted from supernatants using the IndiMag Pathogen Kit (Indical Biosciences), following the manufacturer’s protocol. RNA isolation was performed on the IDEAL™ 32 automated extraction system (Innovative Diagnostics), based on magnetic bead technology.

RNA of PPRV present in the supernatants was quantified using real-time PCR by partial amplification of the N protein gene using the ID GeneTM peste des petits ruminants Duplex kit (Innovative Diagnostics) and then performed with LightCycler (Roche).

### Immunofluorescence staining and inverted microscopy

Cells were fixed at 48 hpi using 4% paraformaldehyde (PFA) for 20 minutes at room temperature. After fixation, permeabilization was performed with 0.1% Triton X-100 in 1X PBS for 5 minutes at room temperature. The cells were then incubated in a buffer composed of 1X PBS, 10% FBS and 1 g BSA per 50 mL for 30 minutes. Primary staining was performed with anti-dsRNA monoclonal antibody J2 (IgG2a, 1 mg/mL, SCICONS™, Jena Bioscience), diluted 1:300, and incubated for 2 hours at room temperature. For detection, cells were incubated with DAPI diluted 1:1000 (Sigma-Aldrich, MBD0015) and either a Donkey anti-Mouse IgG secondary antibody (H+L) Alexa Fluor™ 555 secondary antibody (2 mg/mL) applied at 4 µg/mL (Thermo Fisher Scientific, Cat# A-31570, RRID:AB_2536180), or with TRITC-coupled anti-NPPRV (Mab 38.4) antibody diluted 1:250. In both cases, incubation took place for 1 hour at room temperature, in the dark. The samples were then imaged using an AXIOVERT A1 inverted microscope (Zeiss, France) with Archimed 6.1.4 software, and analysed with ImageJ (version supplied with Java 1.8.0_172 64-bit).

### Statistical analysis

Statistical analyses and histograms were performed using GraphPad Prism v.10.6.1 and R software v.4.5.2. For the analysis of infection rates based on the number of cells infected by the two strains studied on the different hosts, generalized linear mixed models (GLMM) with a binomial distribution were used. Species, cell type, and strain were included as fixed effects, and animal identify and/or experimental batches was included as a random effect to account for repeated measurements within individuals.

Viral titers were analysed using generalized linear mixed-effects models with a negative binomial distribution. Species, cell type and strain were included as fixed effects and animal identify and/or experimental batches as a random effect.

For kinetic experiments, the analysis was performed using generalised mixed models with binomial or beta-binomial distribution in cases of overdispersion, with post-infection time (hpi) as a fixed effect and animal identify and/or experimental batches as a random effect.

RT-qPCR data from kinetics were analysed using linear mixed models (LMM) with the restricted maximum likelihood (REML) method, with hpi as a fixed effect and animal identify as a random effect to account for repeated measurements in the same animal.

Overdispersion and zero-inflation were evaluated where appropriate. Pairwise comparisons between groups were performed using Tukey’s multiple comparison test with adjustment for multiple testing.

## ACKNOWLEDGMENTS

A.Bo., A.Ba. and S.G. are supported by a grant (SI2.756606) from the European Commission Directorate General for Health and Food Safety awarded to the European Union Reference Laboratory for peste des petits ruminants (EURL-PPR, https://eurl-ppr.cirad.fr/).

The authors would like to thank all the staff at UMR ASTRE (CIRAD) for their technical support throughout this study, and in particular Olivier Kwiatek for his help in analysing the titration plates, and Hélène Jourdan for her valuable advice on statistical analyses. We also express our gratitude to the staff of the CIRAD animal facility in Baillarguet — Géraldine Bossard, Thierry Maccotta and Jade Fudaly — for their invaluable assistance. Finally, we would like to thank Aymeric Neyret for his expert advice on immunofluorescence microscopy analyses.

Conceptualisation: A.Bo., V.L., R.-J.E., M.C., P.H., P.T. and A.Ba.; data curation: A.Bo, V.L., M.C. and R.-J.E.; formal analysis: A.Bo., V.L., and P.T.; funding acquisition: A.Ba supervision: A.Ba, P.T. and P.H.; investigation: A.Bo., V.L., M.C. and R.-J.E.; methodology: A.Bo, V.L., R.-J.E, P.T., C.P-G. and S.G.; validation: A.Ba., P.T. and P.H.; visualisation: A.Bo., P.T., P.H. and A.Ba.; writing (original draft preparation): A.Bo.; writing (review and editing): A.Bo, P.H., P.T. and A.Ba.

