## Supplementary material for "Strain-dependent differences in the capacity of peste des petits ruminants virus to infect antigen-presenting cells"

Running Head: APC Infection Differences Among two PPRV Strains

#Address correspondence to Aadel Bouziane (ORCID: https://orcid.org/0009-0007-1090-9110),; address correspondence to Arnaud Bataille (ORCID: https://orcid.org/0000-0002-3508-2144),

*Present address: V. Lasserre, Laboratoire Ecologie et Biologie des Interactions, équipe MHE, UMR CNRS 7267, Université de Poitiers, 5 rue Albert Turpin, 86073 Poitiers Cedex 9, France, Université de Poitiers, Poitiers, France; R.-J. Eloiflin, Institut de Génétique Moléculaire de Montpellier (IGMM), Université de Montpellier, Montpellier, France; P. Holzmuller, Cirad, UMR Selmet, Montpellier, France

The order of authors reflects the relative contributions of the individuals listed up to S. Guendouz. The last three authors, P. Holzmuller, P. Totté, and A. Bataille, contributed to the study in supervisory roles.

KEYWORDS: Morbillivirus, monocyte-derived macrophage, monocyte-derived dendritic cell, virulence, host susceptibility

Supplemental Material

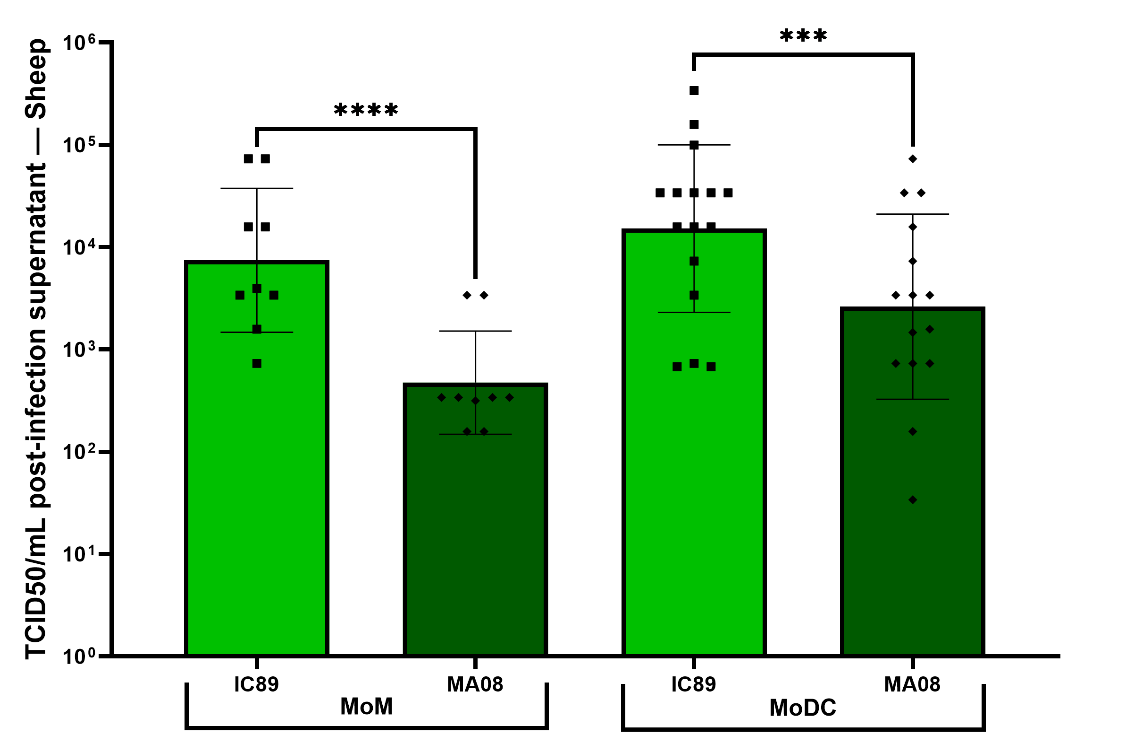

**Supplementary Figure S1: Viral titers (TCID_50_/mL). Comparison of IC89 and MA08 at 48 hpi in sheep MoMs and MoDCs.** Data are shown as individual biological replicates from a minimum of three sheep per condition with mean ± SD; ***P<0.001; ****P<0.0001. Statistical analysis was performed on viral titers using a generalized linear mixed-effects model (GLMM) fitted with a negative binomial distribution. Post-hoc comparisons were performed using estimated marginal means with Tukey adjustment for multiple comparisons.

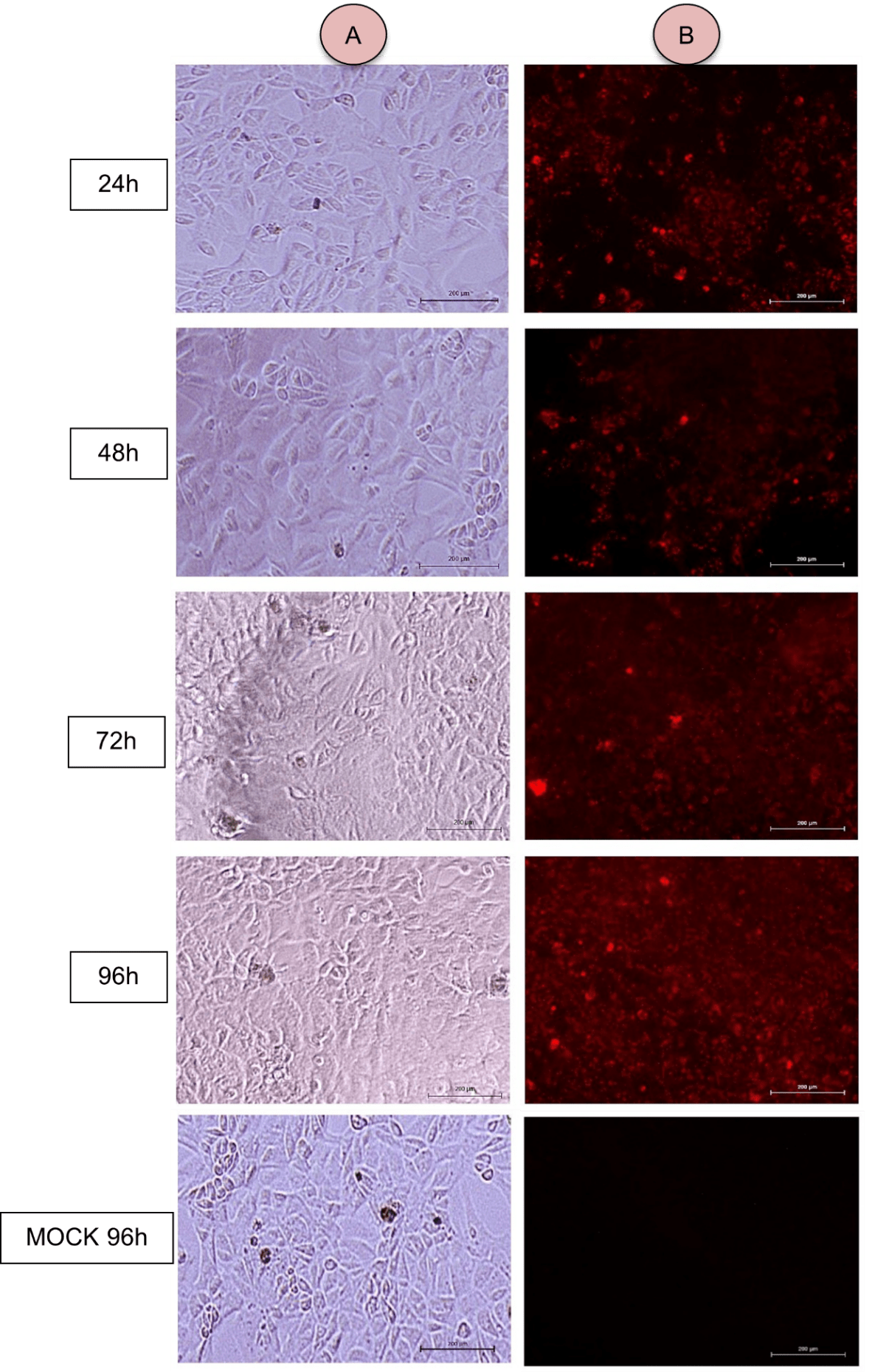

**Supplementary Figure S2:** **Identification of the N protein of PPRV on titration plates of post-infection supernatants from the kinetics of strain IC89 on goat MoMs.** VDS cells were labelled with the 38:4 anti-NPPR antibody coupled to TRITC The titration plate was observed with (B) or without (A) Cy3 filter using the Zeiss AXIOVERT A1 inverted microscope (Zeiss, France) with Archimed 6.1.4 software, magnification x20 (scale: 200 µm).

**Supplementary Figure S3: Immunofluorescent detection of viral double-stranded RNA (dsRNA) in goat MoMs infected with IC89 or MA08.** Infected MoMs were stained with DAPI to identify cell nuclei, dsRNA are detected here in red, scale bar = 25 µm.

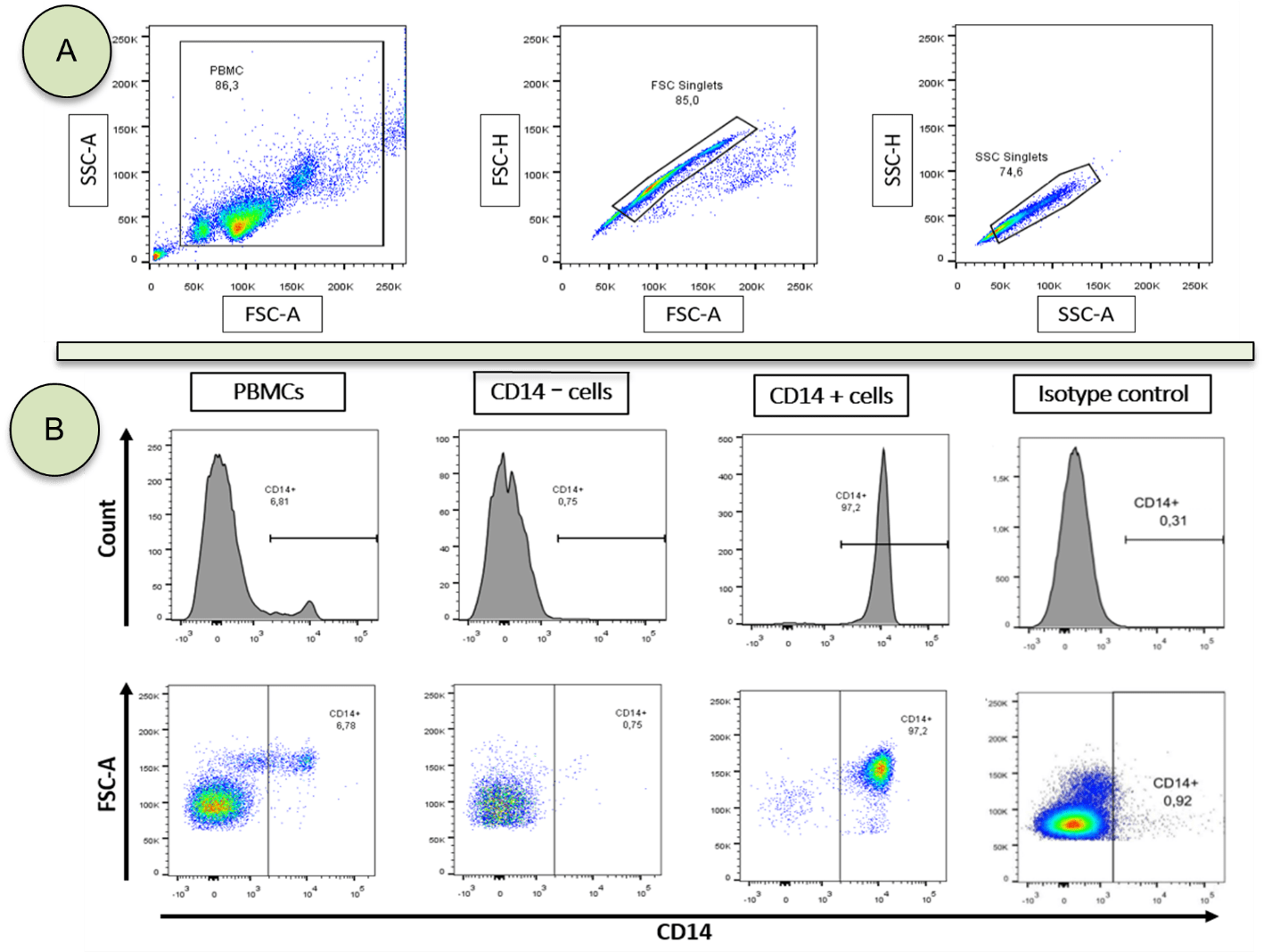

**Supplementary Figure S4:** (A) Gate selection for PBMCs using FSC/SSC, removal of debris and cell doublets using FSC/SSC dot plots. (B) Isolation of CD14+ target cells discriminated by labelling the cells with an IgG1 isotype anti-CD14 antibody by flow cytometry. Example of PBMCs from a goat. The same types of parameters were set for the other goat, sheep and bovine replicates.

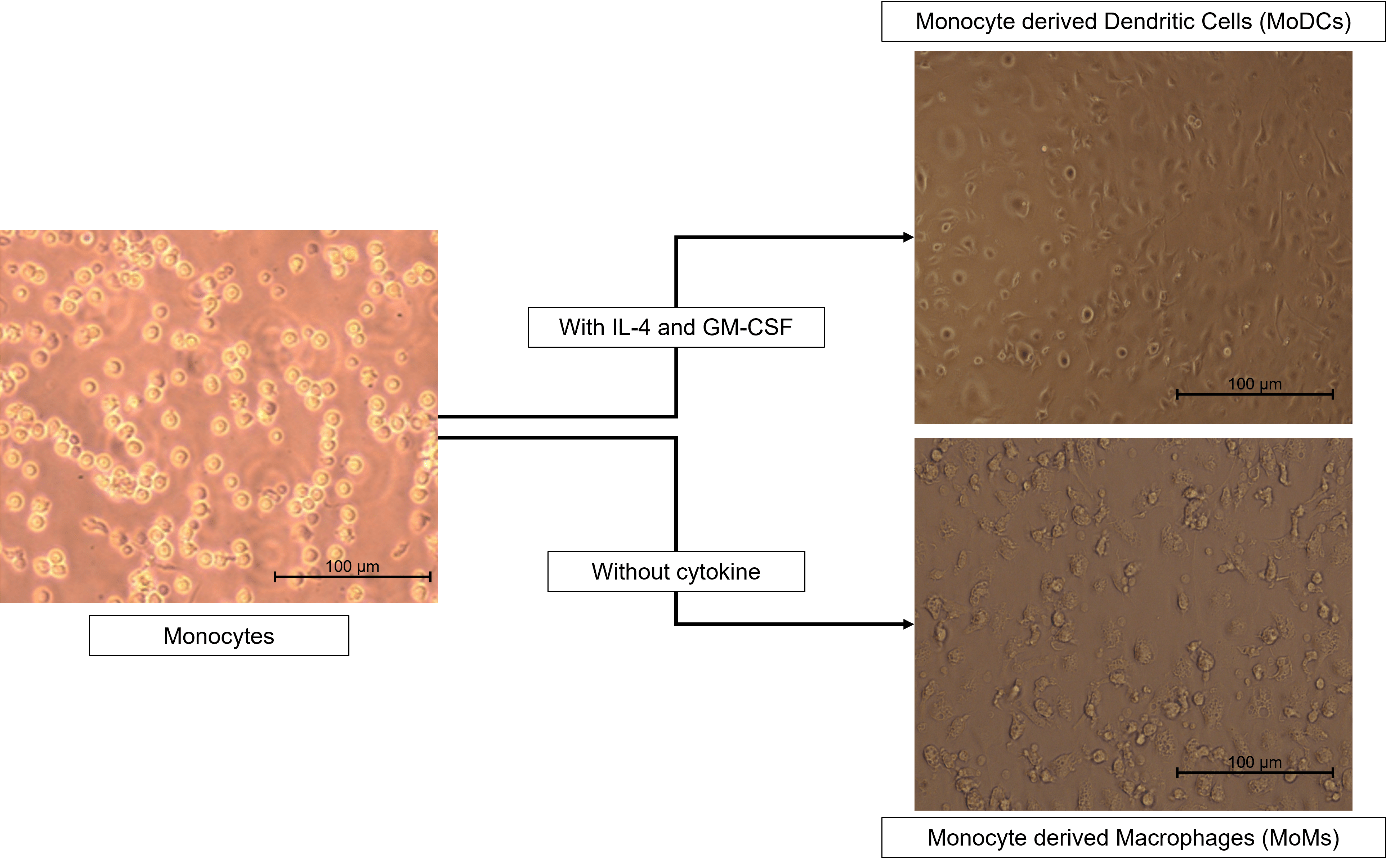

**Supplementary Figure S5: Optical microscope image of monocytes and their differentiation into monocyte-derived dendritic cells (MoDCs) and monocyte-derived macrophages (MoMs).** Adherence and spreading is observed for both MoMs and MoDCS whereas monocytes are round. Dendrites are observed on the surface of rounded MoDCS. This is in agreement with previous work (Park KT et al., 2016; Rodriguez-Martin D et al., 2022; Nfon ck et al., 2012)

(A)

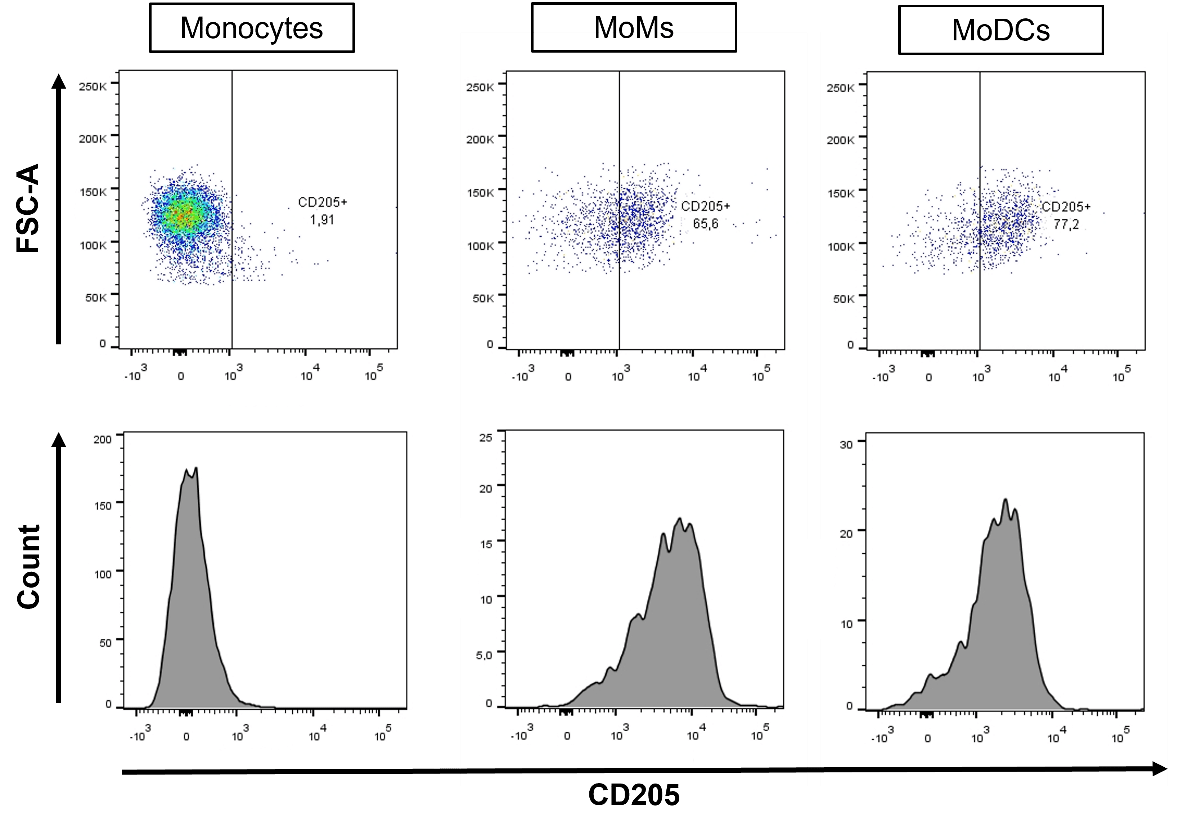

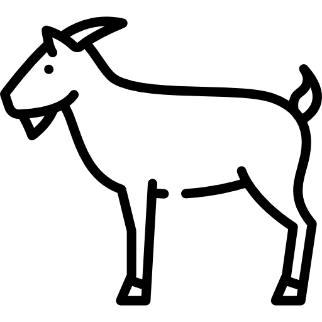

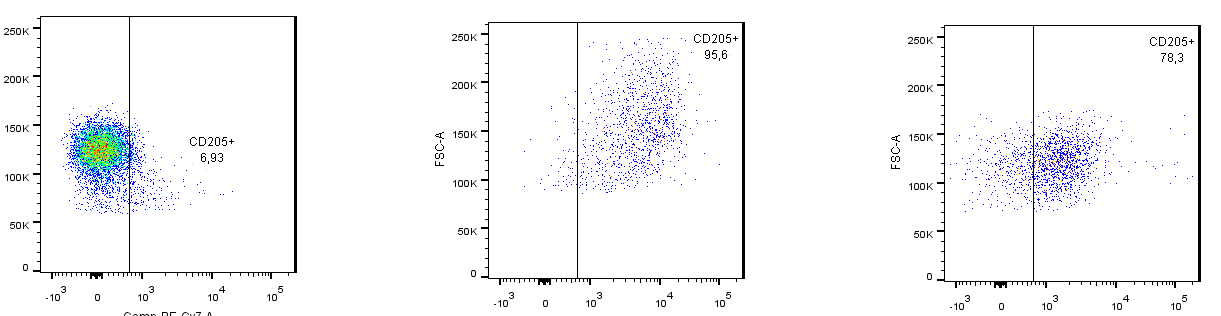

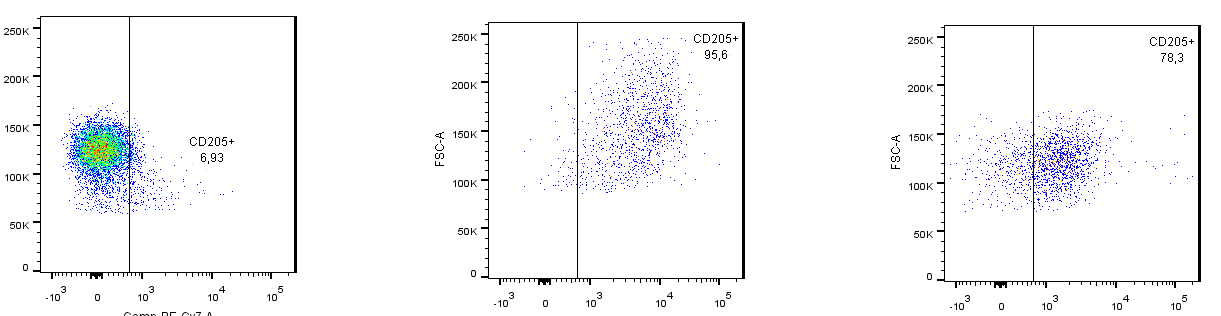

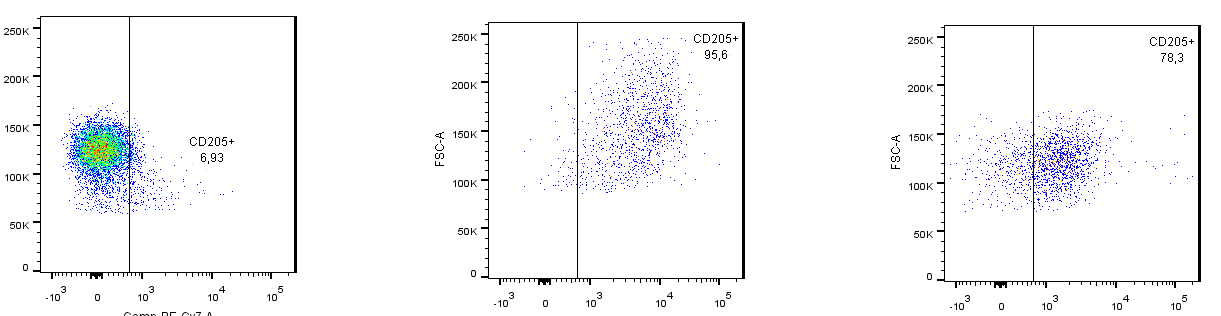

(B)

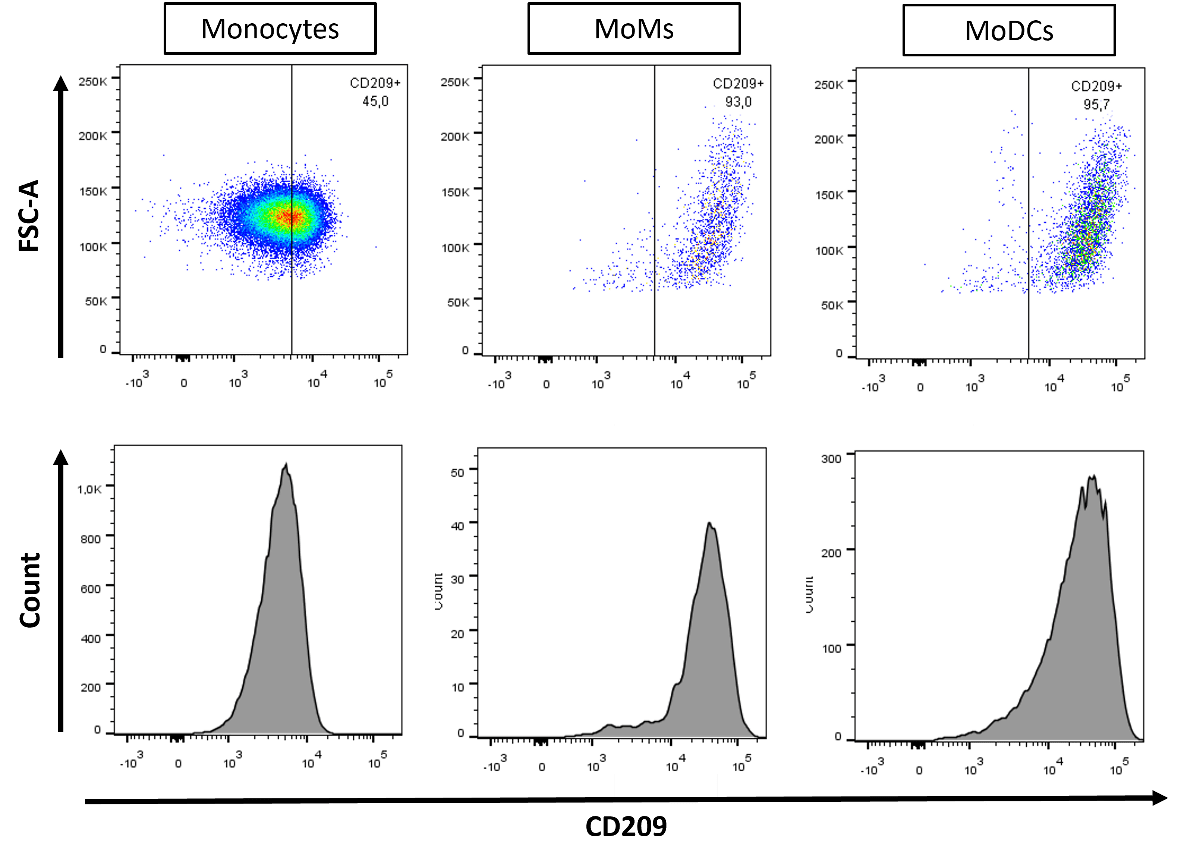

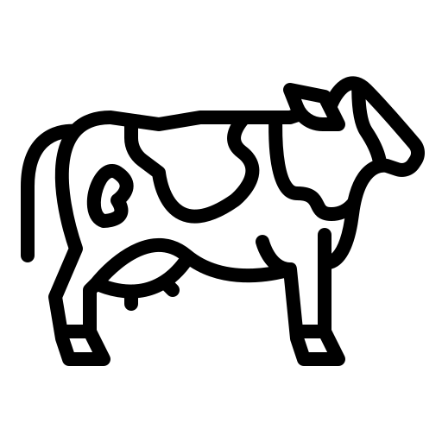

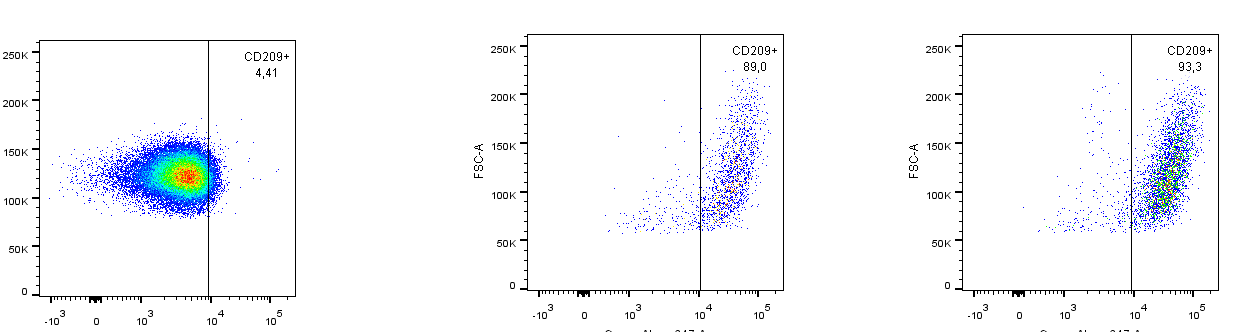

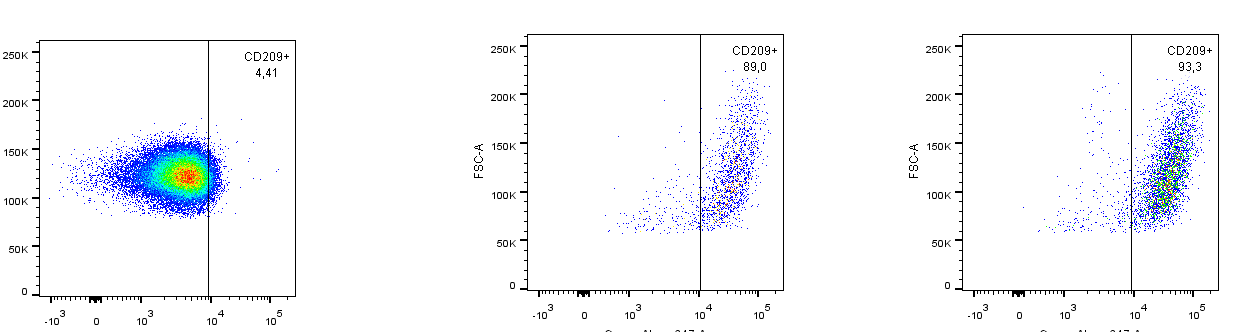

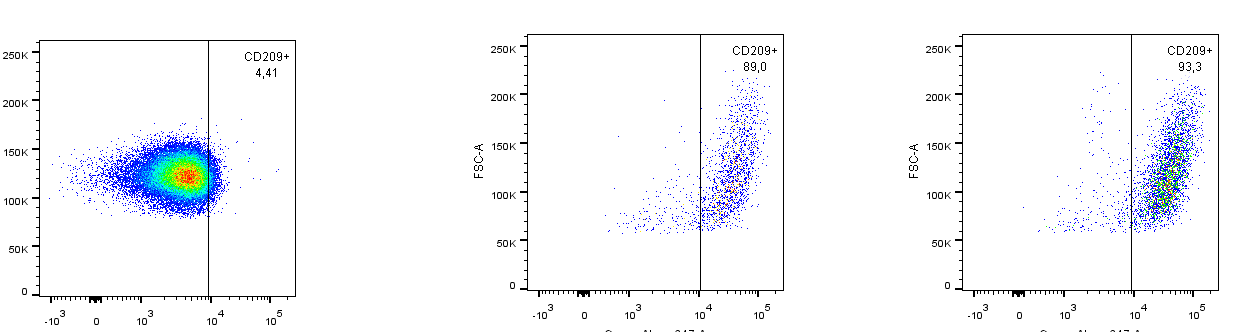

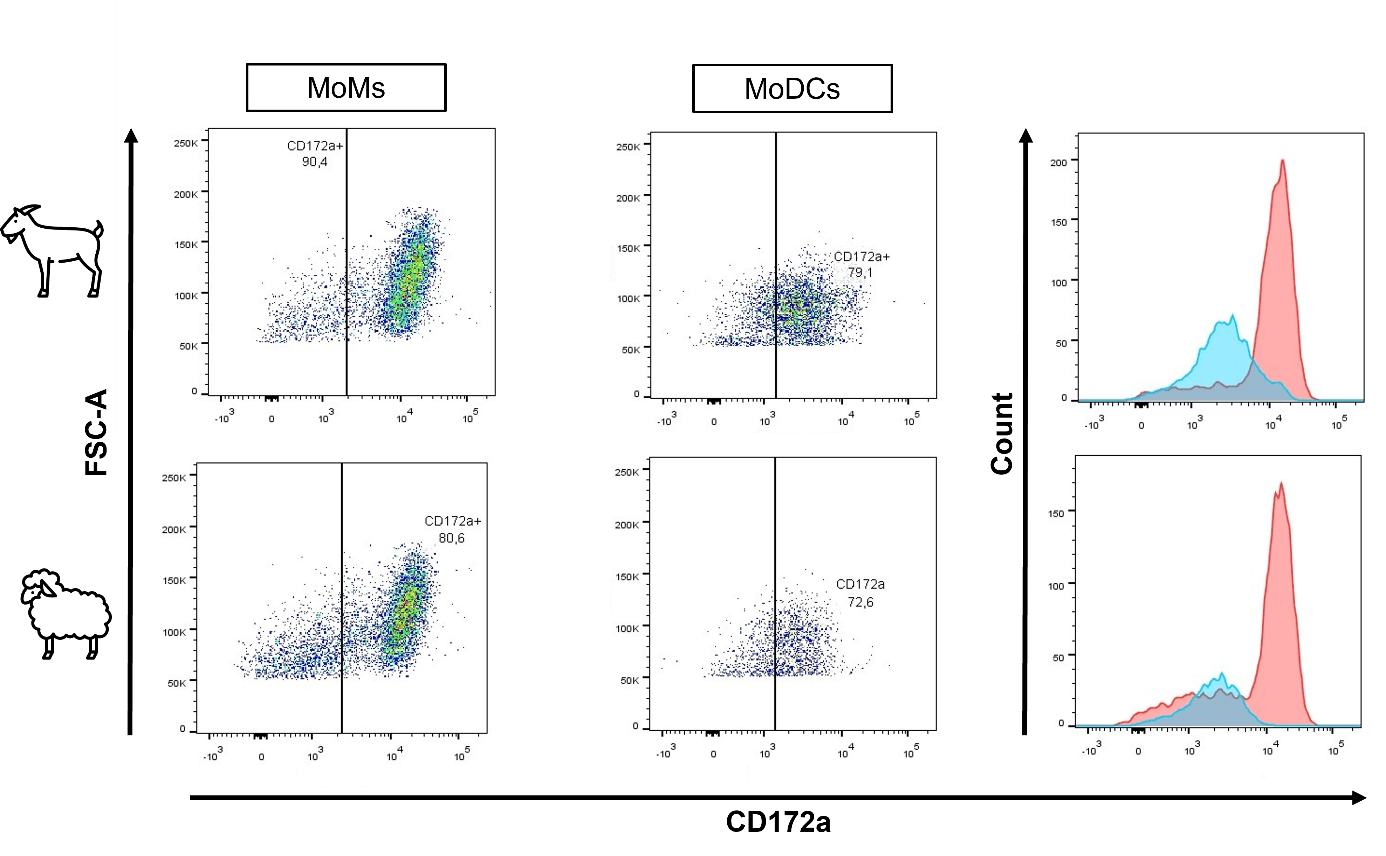

(C)

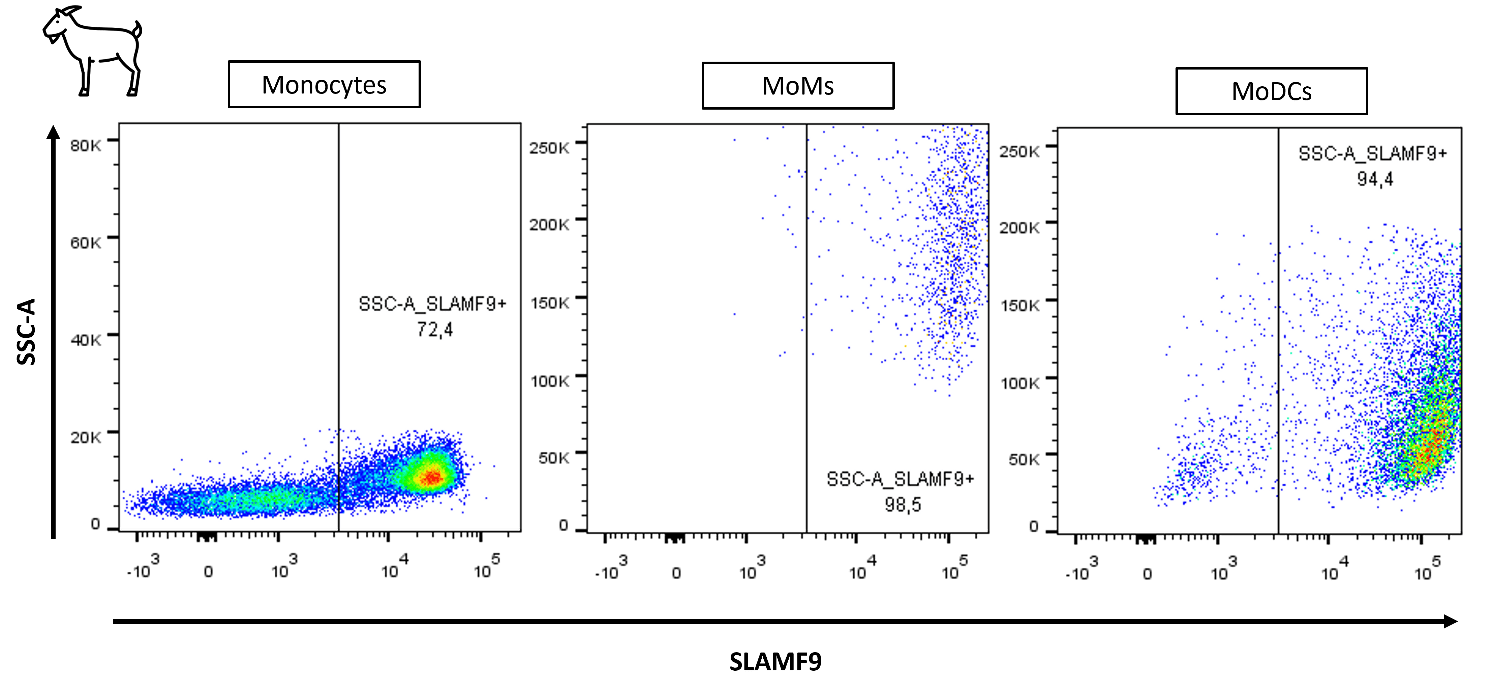

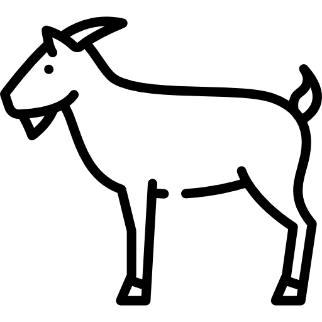

(D)

**Supplementary Figure S6: Phenotypic characterization of MoMs and MoDCs from goat, cattle and sheep.** Representative flow cytometry analysis of (A) goat and (B) cow monocytes, MoMs and MoDCs basedeon (A) CD205 and (B) CD209 expression. Panels show dot plots illustrating (A) CD205 and (B) CD209 expression. Corresponding histograms illustrate the distribution profile of the gated cell populations derived from the dot plots. (C) Representative flow cytometry analysis of MoMs and MoDCs from sheep or goat based on CD172a expression. Histograms show the distribution profiles of the gated populations, with MoDCs represented in blue and MoMs in red.

(D) Representative flow cytometry analysis of goat monocytes, MoMs and MoDCs based on SLAMF9 expression. Panels show dot plots illustrating SLAMF9 expression.

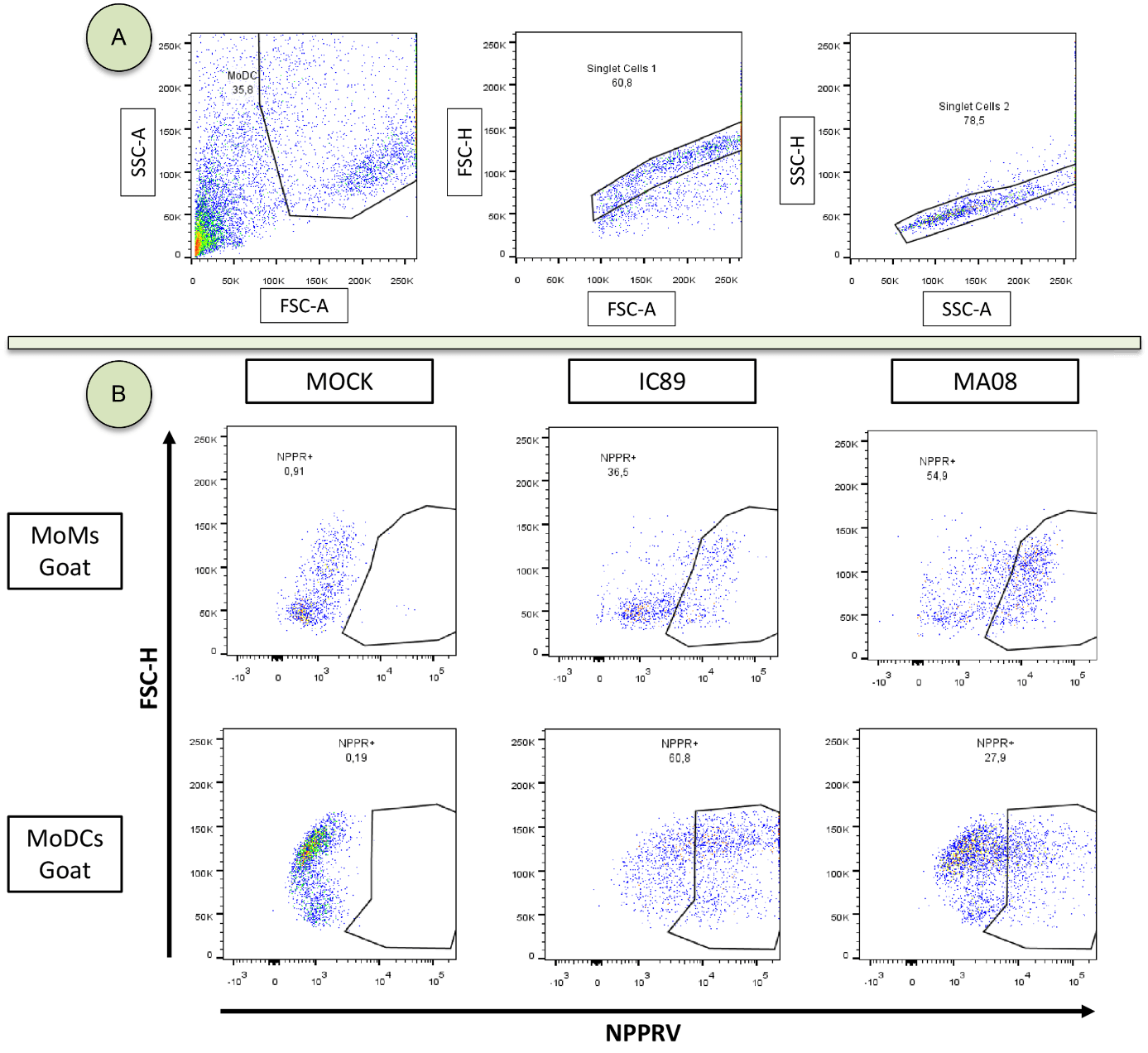

**Supplementary Figure S7:** (A) Gate selection for MoDCs, removal of debris and cell doublets. The same selection was performed on MoMs. (B) Detection of viral nucleoprotein (NPPR+) by flow cytometry on goat MoMs and MoDCs. The same types of parameters have been implemented for other goat, sheep and cattle replicas.
